# Predictive modeling of photoactivation, diffusion and adsorption in cell The case of the optogenetic CRY2-CIBN system

**DOI:** 10.64898/2026.11.19.767830

**Authors:** Antoine Delon, Marc Grosjean, Alexei Grichine, Olivier Destaing, Irène Wang

## Abstract

Strategies combining optogenetics and careful spatiotemporal control of photoactivation are highly promising to decipher the dynamical network of biochemical reactions taking place at the *µ*m scale in living cells. To improve the spatial resolution of such methods, the roles of various molecular processes, such as photochemical reactions, diffusion and interactions, need to be understood and quantified. In this article we develop a minimal and realistic model to describe photoactivation, diffusion and surface-binding of optogenetic probes and validate it thanks to experimental results obtained with a CRY2-CIBN system. A theoretical description of optogenetic activation is implemented, based on the photophysics of CRY2 and on its binding kinetics to the membrane through light-dependent heterodimerization with its partner, CIBN. The cell cytosol is described as a closed reservoir of activable CRY2 molecules. Then, a series of photoactivation pulses of a focused laser beam are applied to activate CRY2 proteins, driving their relocalization to the CIBN molecules anchored at the plasma membrane. We examine both the recruitment rate and the spatial distribution of CRY2 molecules recruited after illumination with either propagative or evanescent light. We have estimated numerical values for the corresponding parameters and put into evidence the key role of the optical saturation of photoactivation. These findings establish a framework to further improve spatial control in optogenetic experiments.

## Introduction

In the mid-2000s [1], biophysical studies of living cells gained a very promising new tool through optogenetics. Thanks to efficient optical techniques, it offered the prospect of triggering intracellular molecular events with high spatial and fast temporal resolutions and therefore, to induce and manipulate out-of-equilibrium reactions in living cells. However, any quantitative assessment of the dynamical network of biochemical reactions taking place within the cytoplasm, nucleus, *µ*m scale organelles, membrane surfaces, etc. implies to properly take into account the underlying chemical and physical processes of diffusion and molecular interactions. This is challenging, as compartmentalisation [2], molecular crowding [3], unknown reaction constants and a huge number of molecular partners make the whole sketch highly complex and difficult to manage [4]. Of particular note is the gradual emergence of whole cell computational models [5], leading to projects such as VCell [6] that aims at simulating cellular structures and functions by reconstructing cellular dynamic behaviors by integrating multiple disciplines, such as physics, biology, computer science and artificial intelligence.

In contrast with the above mentioned numerical strategies, simplified physical models can be used to describe cellular behavior with few parameters and to understand the main processes at play. Here we developed such a tractable biophysical approach of optogenetic recruitment, exemplified with the CRY2-CIBN couple as a system [7]. When photoactivated under blue illumination, the cytosolic CRY2 proteins change conformation and heterodimerize with their CIBN partners, usually anchored at the membrane by a CAAX domain. By fusing CRY2 with a protein of interest, the latter fusion can be relocated and induce vari ous cellular responses at the plasma membrane of cells expressing CIBN-CAAX. Although this heterotypic system has been widely used to drive different cell functions [8-12], its suitability for precise subcellular optogenetic control at the plasma membrane is currently limited by the width of the spatial distribution of recruited proteins, which usual value (≈ 5 µm [13]) far exceeds the area illuminated by the focused laser beam that induces the optogenetic activation. To determine the parameters that govern the spatial resolution of the recruitment at the membrane, we considered the CRY2-CIBN-CAAX system, without fusion to any effector protein. Hence, using this minimal biological system, we studied the following processes: the photoactivation of the freely diffusing CRY2 molecules, their diffusion and anchoring at the membrane. In their seminal article, Valon *et al*. [13] proposed a phenomenological model of photoactivation to describe the recruitment rate at the membrane as a function of the frequency and the number of laser pulses. They assumed that a given fraction of the available CRY2 molecules are activated by each pulse, then diffuse, bind to the membrane through CIBN-CAAX interaction and dissociate after a while, thus refilling the pool of activable molecules in the cytosol. They did not take into account the spatial distribution of the photoactivation laser intensity nor did they consider other dynamic parameters than diffusion. However, the photophysics of the activation process, together with the binding kinetics at the membrane of the activated CRY2 molecules must be taken into account to predict and control the spatial distribution of proteins recruited on the plasma membrane. Moreover, a complete and consistent theoretical model would also make it possible to study the effect of different spatial distributions of laser intensity on the spatial resolution of CRY2 membrane recruitment.

We have recently compared two optical configurations for photoactivation, that both allow to focus light into a spot at the basal membrane of a cell plated on the coverslip on an inverted microscope [14]: a usual propagative laser beam *versus* an evanescent laser beam, obtained by uniformly illuminating the annular region of the back focal plane of a high NA oil objective [15]. The idea was to evaluate to what extent an evanescent spot would induce a sharper localization of recruited CRY2 proteins. Indeed, if diffusion within the cytosol is detrimental to such localization, an evanescent photoactivation should mitigate cytosolic diffusion and thus improve the localization of recruited CRY2 proteins. Ideally, this possibility would make it possible to design CRY2-CIBN optogenetic experiments with spatial resolution reaching the sub-micrometric scale. We observed that the spatial resolution achieved with evanescent illumination, characterized by the Half Width at Half Maximum (HWHM) of the spatial distribution of recruited proteins, was approximately four times wider than the evanescent laser spot itself (≈ 1.4 µm *versus* 0.36 µm). In case of a propagative (i.e. usual) laser spot, the HWHM of the recruited protein distribution was about twelve times that of the measured focused laser spot (≈ 2.8 µm *versus* 0.23 µm, respectively) [14]. Furthermore, it appears that many cells exhibited much wider distributions of recruited molecules.

Thus, we raise the following questions: what are the key parameters governing the spatial distribution of recruited molecules, including the spatial profile and intensity of photoactivation, the photophysical properties, the diffusion and interactions of CRY2 proteins and the geometry of cellular compartments? Can any of these parameters be optimized to enhance resolution? Furthermore, given that the recruitment rate of proteins at the membrane -as excitation pulses repeatedly photoactivate molecules-varies with the illumination mode (evanescent or propagative) [14], we want to investigate its dependence on physical parameters.

These experimental observations and open questions underscore the need for a theoretical model of the underlying molecular processes. The relevance of such a model relies a balance between simplicity (to ensure broad applicability) and predictive power (to ensure practical utility). To our knowledge, existing mathematical descriptions of cellular optogenetics are largely phenomenological or purely descriptive, with only very few grounded in biophysical considerations. Even in the pioneering work of Valon *et al*. [13], and in the recent study by Bohec *et al*. [16], photoactivation is treated as a linear process, without considering the spa-tiotemporal distribution of energy. Additionally, the critical interplay between the intramolecular activation rate of CRY2 and its membrane-binding kinetics is not taken into account. A noticeable exception is the work of Kuznetsov and coworkers [17] who developed a computational framework based on a finite element model. They analysed the signaling dynamics by accounting the complex geometry of cell compartments and the experimental specificities, including the spatiotemporal photoactivation scheme. However, they described the photoactivation process using a constant quantum yield of photoconversion and neglected the depopulation of the ground state. Therefore their approximations are not suited to the present case of a highly focused laser beam. In addition, although their framework can be applied to moderately localized TIRF excitation, they do not report any corresponding experiment. Finally, such numerical simulations are computationally intensive, unlike our simplified model that highlights the physical phenomena.

Regarding spatial resolution, it is worth noting that other optogenetic systems, such as the Phy-PIF photosensitive system, which relies on a two-color control of reversible protein translocation to the membrane [18], could be better suited than CRY2-CIBN to achieve micrometer-scale precision, in spite of longer optical wavelengths. Lov domain based optogenetics has been also used to improve spatial resolution, since the photosensitive system is frequently anchored at the plasma membrane with limited 2D diffusion [19]. Nevertheless, we believe that the global theoretical description we propose here would also benefit to these more complex optogenetic systems.

In the following, we introduce a minimal biophysical model to describe the photoactivation of CRY2 molecules and their membrane recruitment. Then we compare whether this model’s predictions are quantitatively consistent with experimental results obtained from a series of cells expressing the CRY2-CIBN system submitted to propagative and evanescent, tightly focused, illumination modes. We identify the key parameters governing the observed behavior and show that they can be assigned values that render the theoretical description compatible with the experimental results. Finally, in light of this analysis, we discuss possible directions for improving the spatial resolution of membrane recruitment in future optogenetics experiments, especially with the CRY2-CIBN system.

## Results

### Theoretical model

In our biophysical model, represented in Fig. 1A, photoactivation and recruitment of freely diffusing CRY2 proteins occur in a closed cylindrical box of radius *R* and thickness *H*. The CRY2 molecules inside this box, once photoactivated during a laser pulse, inevitably end up binding to the cellular membrane without diffusing or detaching from it on the time scale of the experiment. The first hypothesis of immobile CRY2 molecules on the membrane stems from our observations of recruited protein distributions, the widths of which are stationary [14]. It is also consistent with previous estimations of the lateral diffusion coefficient in the membrane (≈ 0.1 *µ*m^2^/s [13]). The second approximation relies on the fact that the CRY2 residence time at the membrane (≈ 3 min [13]) is longer than the acquisition time (≤ 1 min). This makes an important difference from other works [13, 16, 17], which had to take into account the dissociation kinetics of the CRY2-CIBN dimer to determine the number of recruited proteins. In addition to the dimensions of the closed volume, the relevant parameters of our model are related to the cytosolic diffusion constant of CRY2, *D*, its binding constant at the surface, *K*_*b*_, and the photophysical parameters underlying CRY2 activation. These hypotheses lead to a stepwise recruitment process, as depicted in Fig. 1B: each laser pulse induces an increase in the number of recruited proteins, which stay immobile on the time scale of the experiment (reason why the recruitment curve exhibits plateaus). Correspondingly, the distribution of recruited molecules has a constant width and an increasing amplitude. Finally, because the recruited CR2 molecules originate from the cytosol (and do not get back to), the recruitment rate slows down as the cytosol empties.

**Figure 1.**
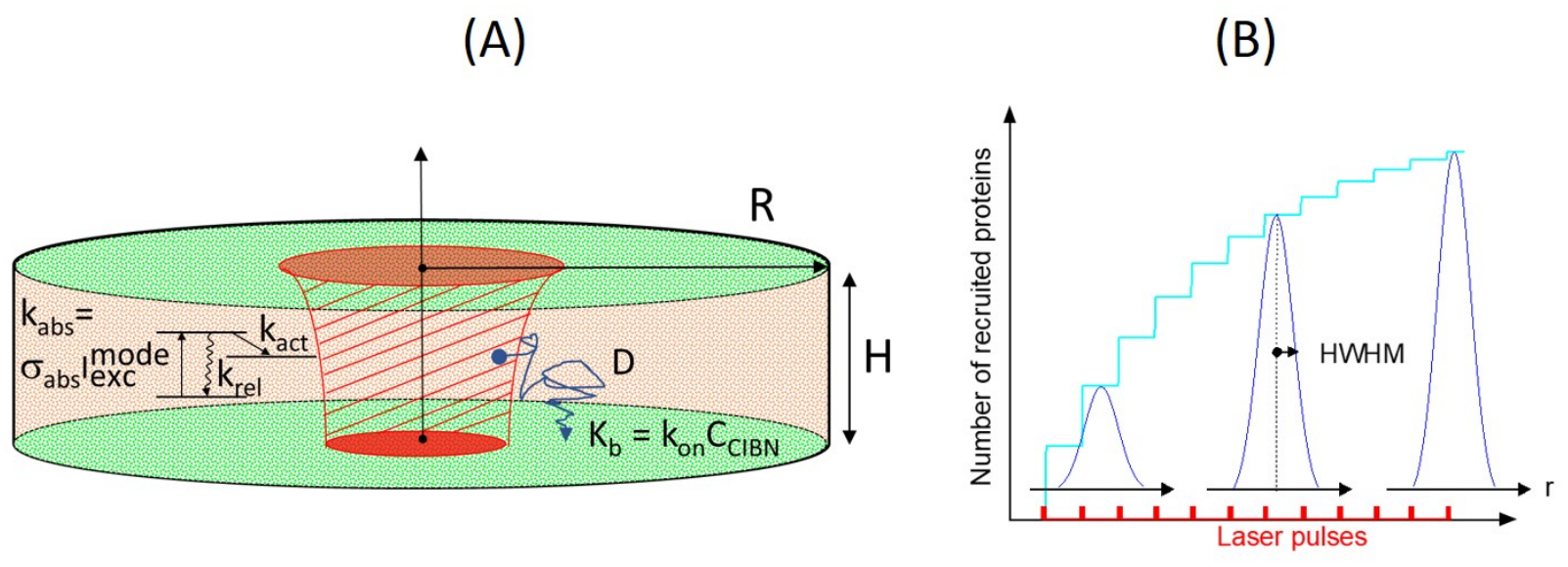
Cell model and recruitment. (A) The cell is represented as a flat cylinder, of radius *R* and thickness *H*, containing a uniform concentration of CRY2 molecules. They can be photoactivated by the incoming laser beam (inner zone shaded in red), according to the photophysical sketch shown on the left. Molecular diffusion (blue random trajectory) leads to irreversible binding of photoactivated CRY2 molecules with CIBN molecules (greenish) that cover the basal, apical surfaces and the lateral wall. (B) Each laser pulse induces a step in the number of recruited proteins that stay immobile on the surfaces. The corresponding surface distribution increases in amplitude, while its width remains constant, as illustrated by the radial distribution plotted at different time steps. Finally, due to the emptying of the cytosolic reservoir of CRY2 molecules, the recruitment rate progressively slows down.

As detailed in section S1 of Supporting Information, the theoretical derivation starts by calculating the number and the spatial distribution of photoactivated CRY2 molecules just after one laser pulse. This is obtained with the simplest consistent scheme, a three-level system, and involves the absorption cross section *σ*_a_ , the pulse duration, *δt*_pulse_, and the spatial distribution of the laser intensity, 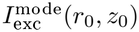 , the relaxation and activation rates, *k*_rel_ and *k*_act_. In another step, considering an activated molecule at a given initial position (*r*_0_,*θ*_0_,*z*_0_), we computed its 3D distribution at time *t* taking into account its diffusion and surface binding properties. Then, the molecular flow at the surface is integrated from *t* = 0 to ∞ to yield the final distribution of molecules recruited at the surface. To obtain the total distribution of recruited CRY2 proteins, we integrate over all initial positions weighted by the 3D distribution of photoactivated molecules, calculated in the first step.

As explained in the subsection *Quantitative comparison of theoretical and experimental results* - *Prior assessments*, many parameters have been assigned values from the literature, so that the only free remaining parameters are the activation rate, *k*_act_ and the binding constant at the membrane, *K*_*b*_.

We now discuss their influence on the recruitment rate and on the width of the distribution of molecules recruited at the basal surface.

#### The slowdown of the recruitment rate gets more pronounced as the activation rate *k*_act_ in-creases

In the section S1 of Supporting Information, we show that the slowdown of the recruitment rate at the membrane depends solely on the intramolecular activation rate *k*_act_. Indeed, in our model, the relative number of recruited CRY2 molecules only depends on the average, over the cell volume, of the fraction of activated CRY2, 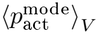 , since any activated molecule will ultimately and irreversibly bind to the membrane, whatever the *K*_*b*_ binding constant. Fig. 2 shows that the slowdown of the recruitment rate becomes more pronounced when *k*_act_ increases, the latter governing the fraction of activated CRY2, and hence the relative depletion of cytosolic CRY2 induced by each laser pulse. In addition, the slowdown of the recruitment rate is more important in the propagative illumination mode.

**Figure 2.**
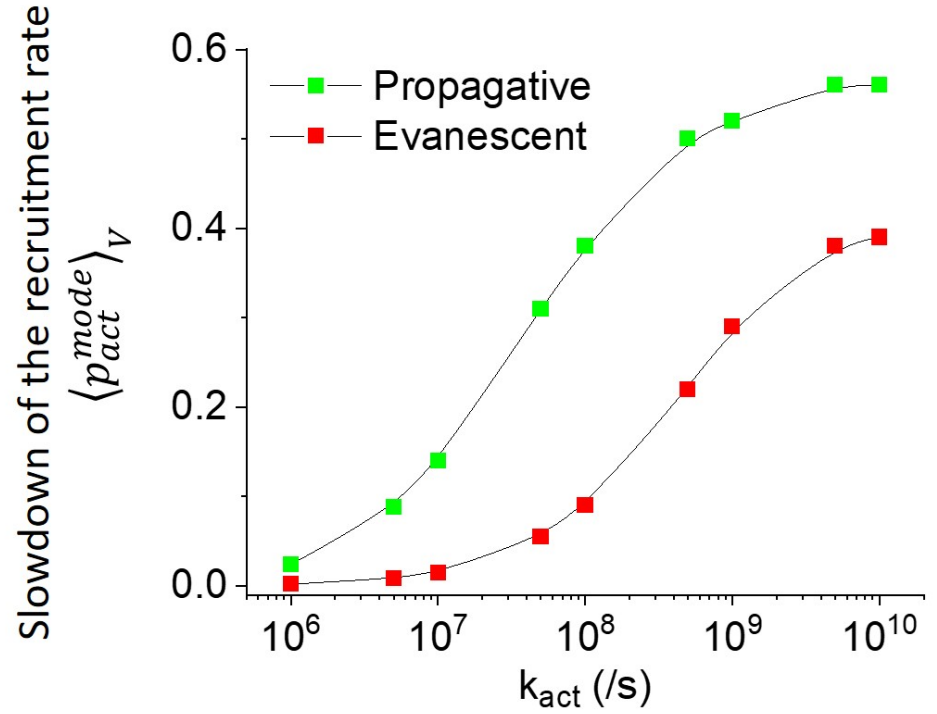
Effect of the intramolecular activation rate, *k*_act_, on the slowdown of the recruitment rate for the two illumination modes, propagative and evanescent. This quantity corresponds to the average, over the cell volume, of the fraction of activated CRY2 molecules, 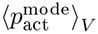.

#### The width of the distribution of recruited molecules depends on both activation rate *k*_act_ and binding constant *K*_*b*_

The width of the distribution of molecules recruited on the membrane depends on both the activation rate, *k*_act_ and the binding constant, *K*_*b*_. On one hand the *k*_act_ rate increases the photophysical saturation of the activation process, thus widening the spatial distribution of the activated population beyond that of the illumination intensity. On the other hand, the larger the binding constant, *K*_*b*_, the narrower the distribution of molecules recruited at the membrane: indeed, as *K*_*b*_ increases, the activated molecules are less likely to diffuse back and forth between the surfaces. Henceforth, they are captured more rapidly upon reaching a membrane boundary, reducing the distance over which they diffuse, thus narrowing the membrane recruitment profile. Fig. 3 shows, for both illumination modes, the counterbalancing effects of *k*_act_ and *K*_*b*_ on the width of the distribution of recruited molecules.

**Figure 3.**
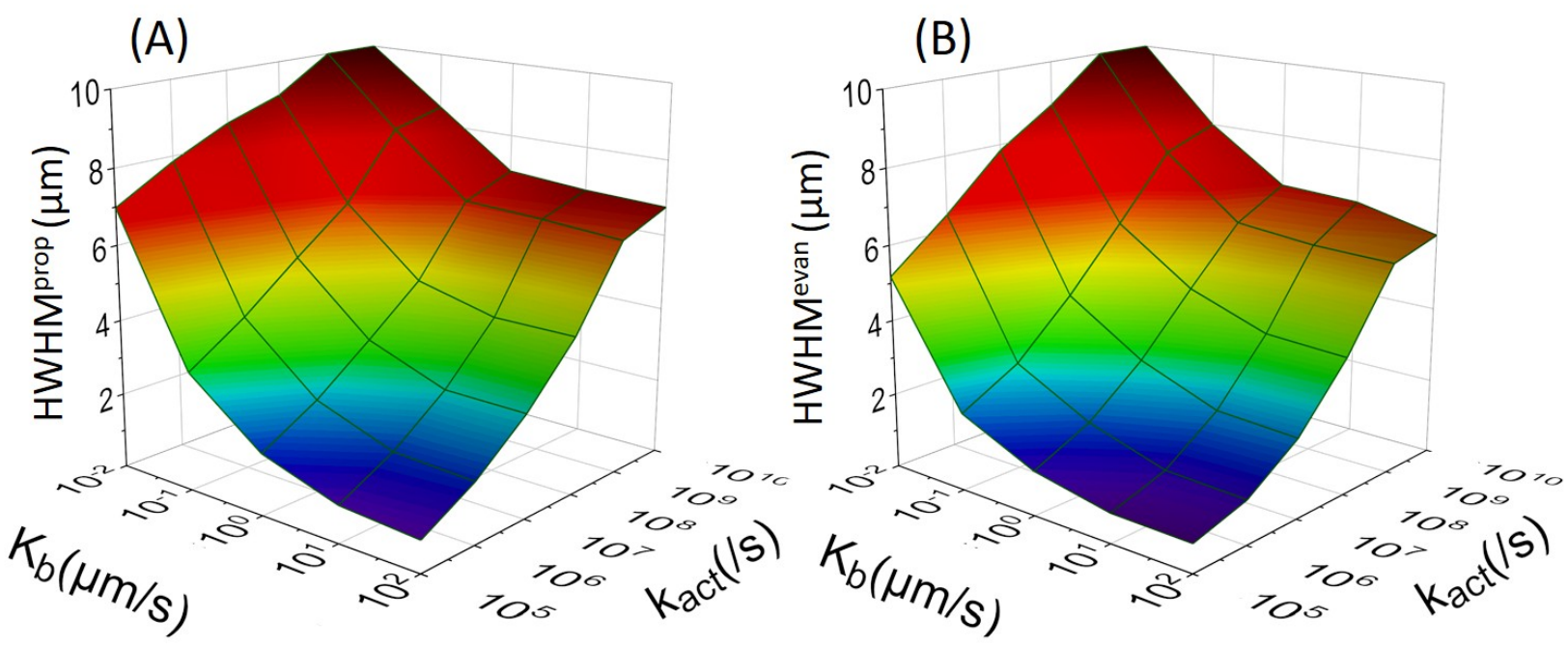
Effect of the activation rate, *k*_act_ and of the binding constant, *K*_*b*_, on the half width at half maximum (HWHM) of the distributions of recruited molecules, for the propagative (A) and evanescent (B) illumination modes. Due to the photophysical saturation of the activated population, an increase in *k*_act_ tends to widen the distributions, while an increase in *K*_*b*_ forces the molecules to bind the surfaces whilst minimising diffusion.

### Experimental results

Examples of 561 nm-excited TIRF images of CRY2-mCherry recruited to the basal membrane are depicted in Fig. 4, which shows one cell of each batch (batch-1 and batch-2).

**Figure 4.**
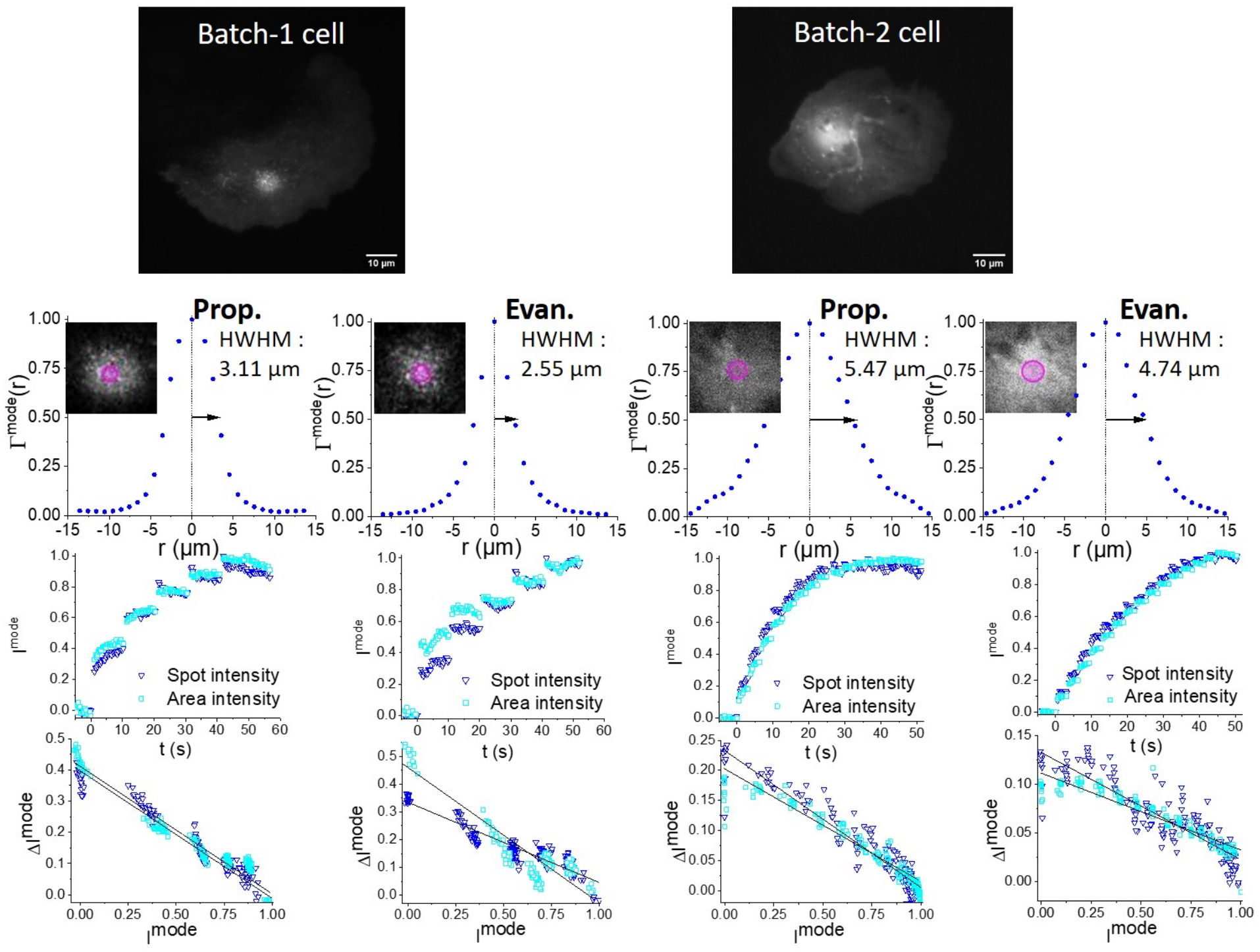
Examples of experiments performed on two cells, from the batch-1 (left-hand side) and from the batch-2 (right-hand side). For each case, the averaged mCherry fluorescence image is shown at the very top (scale bars are 10 *µ*m long). The radial distributions of recruited proteins at maximum recruitment, Γ^mode^ (*r*), are shown just below (1^*st*^ line of graphs), for the propagative and evanescent modes, with zooms on recruitment spots as insets (superimposed circles are 1 *µ*m in radius). The bottom part of the figure displays, in the 2^*nd*^ line of graphs, the fluorescence intensity, *I*^mode^, either measured at the laser spot (Spot intensity, triangles) or integrated over the cell area (Area intensity, squares), along the series of photoactivation pulses. Finally, at the bottom (3^*rd*^ line of graphs), the intensity steps induced by the successive laser pulses, Δ*I*^mode^, are plotted *versus* the intensities before each pulse (to simplify writing, *I*^mode^ and Δ*I*^mode^ notations do not show *k* superscripts). Black lines correspond to linear fits.

The two features extracted from these images are the radial distribution of recruited proteins, quantified by the HWHM of the bright region when the recruitment is maximal (1^*st*^ line of graphs underneath the two top images in Fig. 4), and the evolution of the intensity at the laser spot (Spot intensity) or of the intensity integrated over the cell area (Area intensity) as a function of time (2^*nd*^ line of graphs in Fig. 4). See the *Methods* section for details about the estimations of these experimental outputs.

These measurements may be influenced by 3 variables or factors: the excitation pulse duration (0.05, 0.1 and 0.5 s), the state of the cells (batch-1 or 2) and the illumination mode (propagative or evanescent). To determine rationally whether our data depend on those factors, we performed a Generalized Linear Model (GLM) [20], using the slowdown of the recruitment rate or the distribution width as the dependent variable and the 3 above-mentionned factors as explanatory variables. We will not report all the results of GLM analysis, but its outputs will guide us in the following discussion.

#### Slowdown of the recruitment rate

As detailed in the section *Methods* - *Analysis of molecule recruitment rate and distribution width*, the fluorescence intensity curve as a function of time can be converted into a graph showing the intensity step, Δ*I*^mode^, as a function of the intensity, *I*^mode^. Examples of such graphs, inferred from the experimental data, are shown on the bottom line of Fig. 4. In agreement with the derivation presented in section S1 of Supporting Information (Eq. 41), we verified experimentally that Δ*I*^mode^ depends linearly on *I*^mode^. This simply means that the number of photoactivated proteins after each pulse (which leads to the intensity increase Δ*I*^mode^) is proportional to the concentration of remaining CRY2 in the cytosol, which decreases linearly with the number of proteins already bound to CIBN-CAAX at the membrane (proportional to *I*^mode^), since the total number of CRY2 is unchanged. The number of recruited CRY2 approaches a saturation plateau over time. The corresponding slowdown parameter, *B*^mode^, can be directly compared to the relative population of activated proteins derived theoretically, noted 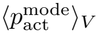 (see Eq. 36 to 41 in section S1 of Supporting Information).

One of the main results of the GLM analysis is the dependence of the recruitment slowdown parameter upon the illumination mode: the slowdown of the recruitment rate is more pronounced in propagative mode compared to evanescent mode, *B*^prop^ *> B*^evan^, as shown on Fig. 5. This is expected since evanescent waves are confined in a thin layer close to the cell membrane, so that the overall number of proteins that can be photoactivated in this case is lower, as compared to a beam propagating through the whole thickness of the cell (even if the intensity of the evanescent wave is enhanced at the interface).

**Figure 5.**
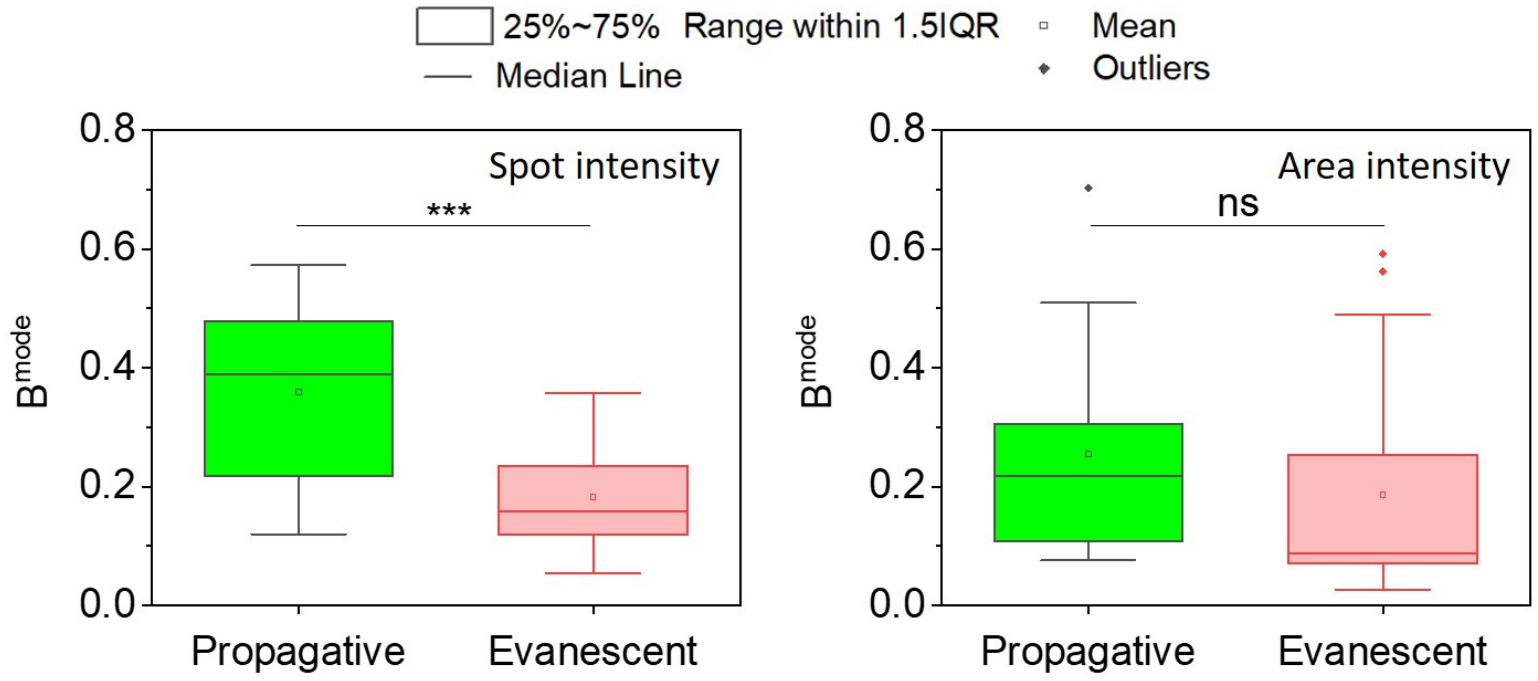
Slowdown of the recruitment rate of propagative and evanescent illuminations, for the two analysis methods, Spot intensities (left) and Area intensities (right) (different *δt*_pulse_ have been grouped in these data). *p*-values from ANOVA analysis are represented: ^***^ stands for *p <*0.001 and ns for nonsignificant.

In Table 1, the values (median, minimum and maximum) of *B*^mode^ are presented for different pulse durations, *δt*_pulse_ Although the limited statistics of our data calls for caution, we observe that the recruitment slowdown parameter tends to increase with the pulse duration from 0.05 to 0.5 s. This is also expected since the longer the excitation pulse the more proteins can be photoactivated, as shown by Eq. 2 in section S1 of Supporting Information.

**Table 1:**
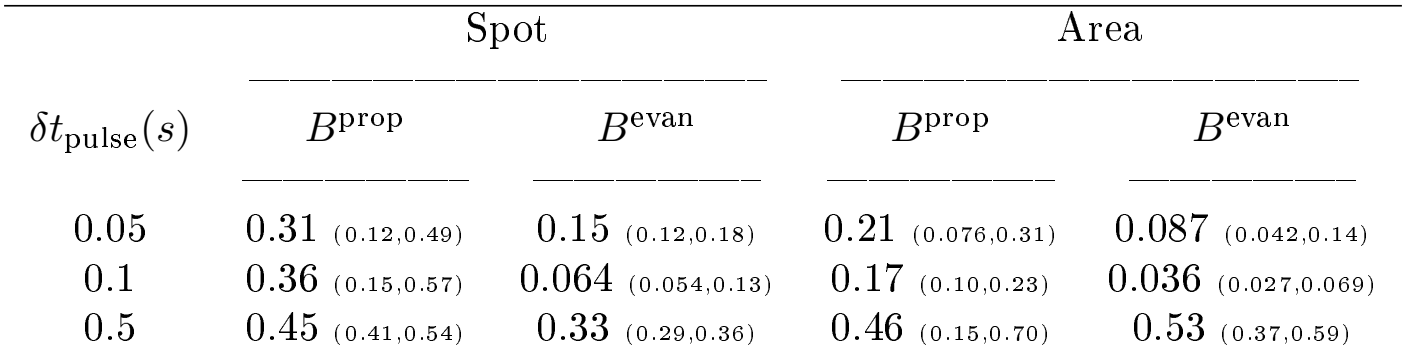
Medians, with minimum and maximum values in parenthesis, of the experimental recruitment slowdown parameter, *B*^mode^, for the different acquisition parameters (propagative, evanescent modes and pulse duration, *δt*_pulse_) and the two analysis methods (spot and area integrated intensities).

|  | Spot |  | Area |  |
|---|---|---|---|---|
| *δt*<sub>pulse</sub>(*s*) | *B*<sup>prop</sup> | *B*<sup>evan</sup> | *B*<sup>prop</sup> | *B*<sup>evan</sup> |
| 0.05 | 0.31 (0.12,0.49) | 0.15 (0.12,0.18) | 0.21 (0.076,0.31) | 0.087 (0.042,0.14) |
| 0.1 | 0.36 (0.15,0.57) | 0.064 (0.054,0.13) | 0.17 (0.10,0.23) | 0.036 (0.027,0.069) |
| 0.5 | 0.45 (0.41,0.54) | 0.33 (0.29,0.36) | 0.46 (0.15,0.70) | 0.53 (0.37,0.59) |

Additionally, the GLM analysis showed that the cell batch has no influence on *B*^mode^ , which is also in agreement with our model. Indeed, this quantity is only linked to the photoactivation of CRY2 which is governed by photophysical parameters at the molecular level. These intramolecular parameters (*σ*_abs_, *k*_rel_ and *k*_act_) should depend only very slightly on the state of cells in the different batches, contrary to the binding constant at the membrane, *K*_*b*_, that is related to cell-dependent parameters such as the CAAX-CIBN protein expression level.

#### Spatial distribution of recruited proteins

According to the GLM analysis, the width of the distribution of recruited molecules depends mainly on the cell state and the illumination mode (*p* ≈ 0.1 for these 2 explanatory variables), while the pulse duration has a negligible influence (*p* ≈ 0.3).

The impact of the cell state on this width is dramatic, as shown on Fig. 6: the median (minimum, maximum) value of HWHM^mode^ goes from 2.6 *µ*m (2, 3.5) for batch-1 cells to 5.7 *µ*m (3.7, 9.3) for batch-2. While the origin of such a strong disparity is unclear, it may result from variations in the surface density or accessibility of CIBN between cell batches. Indeed, our model showed that the recruitment width depends on both the activation rate, *k*_act_ and the binding parameter, *K*_*b*_ which is the product of the CRY2-CIBN bimolecular reaction rate with the surface density of CIBN molecules. The density of CIBN molecules on the cell membrane may vary between cells of different batches. Moreover, differences in membrane organization, cytoskeletal state or local molecular crowding, can also affect the encounter and binding of activated CRY2 to membrane-anchored CIBN.

**Figure 6.**
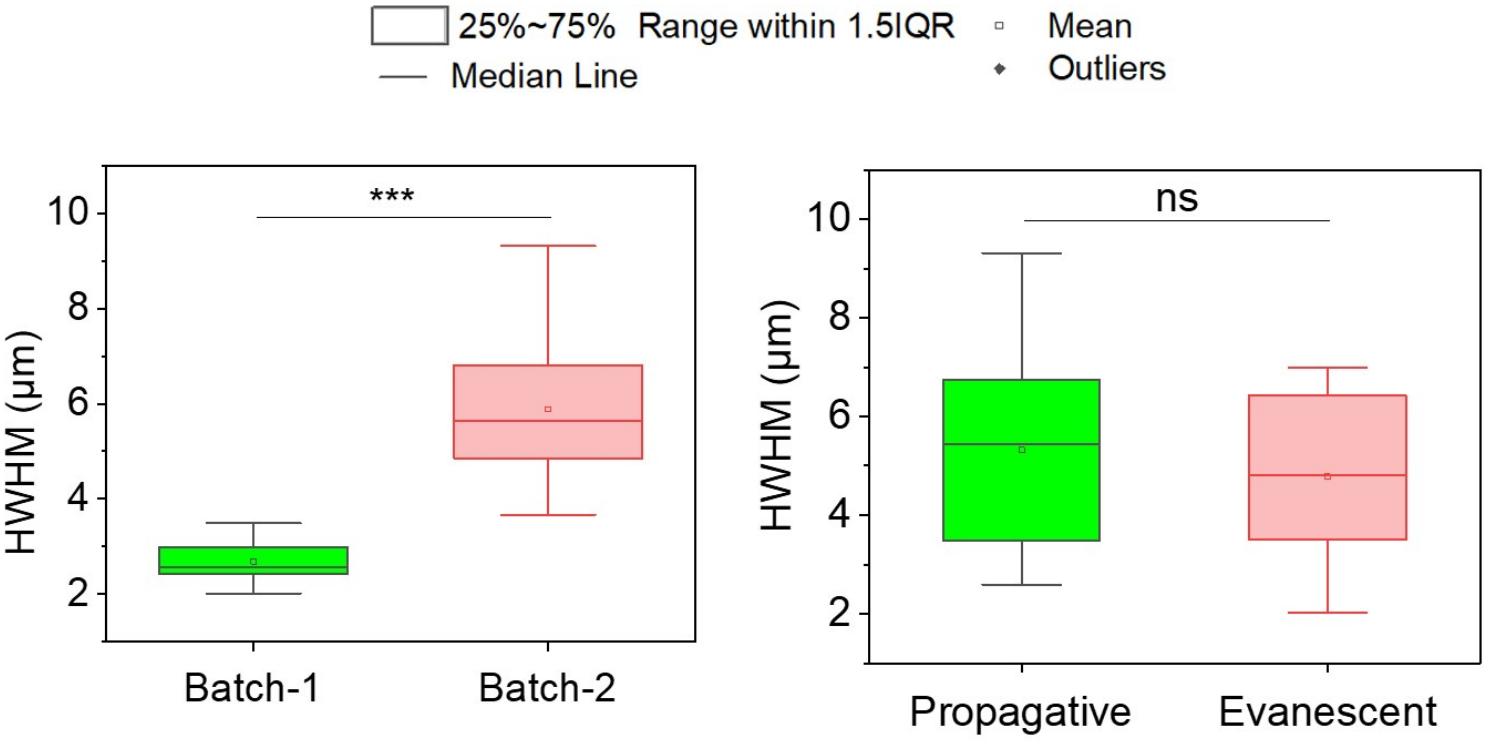
HWHM of the membrane distribution of recruited proteins for the two cell batches (left) and the two illumination modes (right). *p*-values provided by an ANOVA analysis are given, ^***^ standing for *p <*0.001 and ns for nonsignificant.

Whether the HWHM depends on the illumination mode is not clear, as shown in Fig. 6. To refine the analysis, we paired the widths found for propagative and evanescent illuminations on the same cell. On Fig. 7, one can see that evanescent illumination generally yields a narrower distribution than a propagative one, since most data points are in the HWHM^evan^ *<* HWHM^prop^ zone. A Student’s paired t-test, where the hypothesis alternative to the null one is HWHM^evan^ *<* HWHM^prop^, gives a *p*-value of 0.092. Consistently, the model also predicts a slightly narrower distribution in the evanescent case, even if our experimental conditions seem characterized by a strong saturation of the photoactivation, resulting in distributions of molecules much wider than the laser focus, whatever the illumination modes.

**Figure 7.**
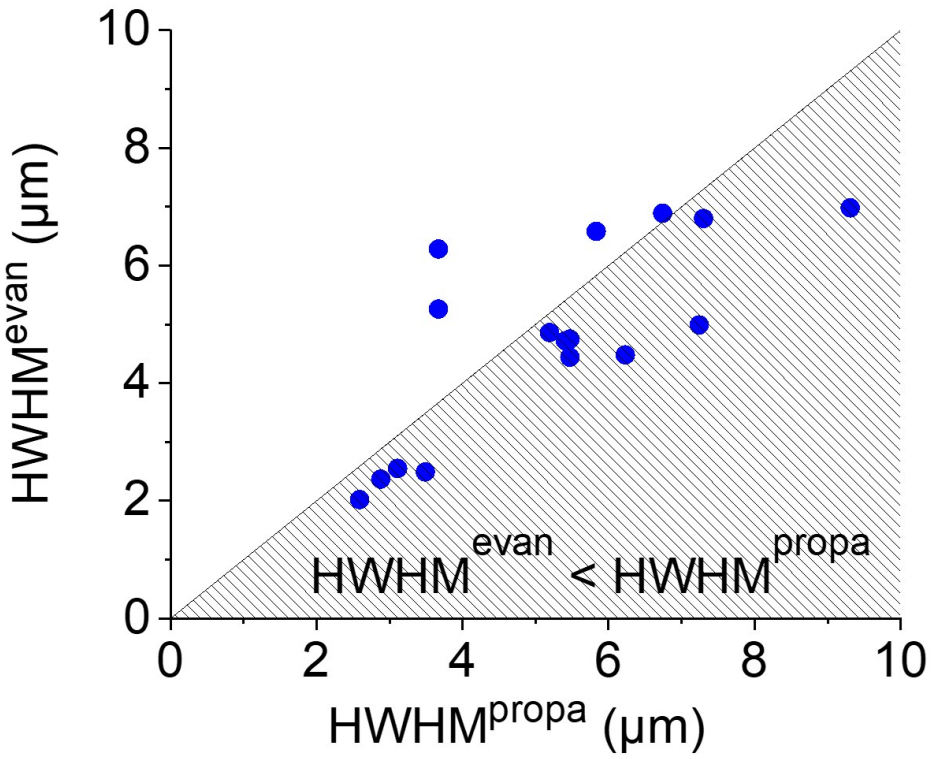
HWHM of the distributions of recruited proteins paired by cell: each HWHM^evan^ value is represented versus the HWHM^prop^ value for the same cell. A majority of data points are within the shaded zone, corresponding to HWHM^evan^ *<* HWHM^prop^.

### Quantitative comparison of theoretical and experimental results

We now aim to quantitatively describe the experimental results using our model with a set of reasonable parameters.

#### Prior assessments of parameters

First, several parameters are fixed based on previous literature. The absorption cross section, *σ*_a_ , is set to 10^*−*16^cm^2^, a typical value at the maximum of the absorption curve of an allowed electronic transition, which also agrees with previous estimations for CRY3 [21]. After excitation, the molecular fate is governed by the relaxation rate to the ground level, *k*_rel_ , and the activation rate to the photoactivated level, *k*_act_. Although they play independent roles, according to Eq. 3 to 6 in section S1 of Supporting Information, our experimental data is insufficient to determine both of them. Therefore, we have decided to fix *k*_rel_ at a reasonable value of 10^9^ s^*−*1^, consistent with the literature [21] and use *k*_act_ as a free parameter to adjust the model to experimental results. In our images, we observed that photoactivated CRY2 tend to oligomerize and form clusters, which is a known reaction [12, 22]. Therefore, we set the diffusion constant to 1 *µ*m^2^/s, which is ten times slower than the value usually reported [12, 13].

The parameter governing the CRY2 binding constant at the surface is *K*_*b*_ = *k*_on_*C*_CIBN_, where *k*_on_ is the bimolecular reaction rate between CRY2 and CIBN and *C*_CIBN_, the surface density of anchored CIBN molecules. Note that we neglect any saturation of the number of available CIBN sites upon adsorption, in agreement with [13]. To determine the interval of possible values for *K*_*b*_, we first consider *k*_on_. According to Ref. [23], the rate of many protein-protein interactions are in the 10^4^ ∼ 10^6^ /M/s range, i.e. *k*_on_ is in the 2 × 10^*−*5^ to 2 × 10^*−*3^ µm^3^/s range. Interestingly, using a CRY2 residence time of ≈ 3 min [13], this corresponds to a membrane binding affinity (or equilibrium dissociation constant *K*_*d*_) in the 6 ∼ 600 nM range. This is consistent with the general consideration that light-sensitive protein affinity is generally in the range of a few to a few hundred nM [24, 25]. That said, an upper limit of *C*_CIBN_ can be estimated by considering the case of proteins densely packed on the membrane, with a mean distance of 5 nm. Altogether, a conservative range of possible values for *K*_*b*_ would be 0.1 to 100 *µ*m/s.

The aim is now to adjust the activation rate of CRY2, *k*_act_, and the binding constant, *K*_*b*_ (within its expected range), using experimental results to check the consistency of our model. We expect *k*_act_ to be mostly constant across cells, whereas *K*_*b*_ may vary from one cell to another since it depends on the density of CIBN on the membrane.

#### Constraints imposed by the recruitment slowdown parameter on *k*_**act**_

Let us first consider the recruitment slowdown parameter: as previously explained, the experimental values (*B*^mode^) can be compared directly to the average, over the volume of the cell, of the relative population of activated CRY2 proteins, 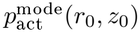, given by Eq. 2 in the section S1 of Supporting Information. The theoretical values, shown in Table 2, increase with *k*_act_ as expected. If we choose a cell volume of radius *R* = 10 µm and height *H* = 4 µm (which seems close to the dimensions of the cells in experiments), we find that, to fit the experimental slowdown parameter (see Table 1), the photoactivation rate *k*_act_ must be higher than 10^8^ /s.

**Table 2:** Theoretical values of the recruitment slowdown parameter, 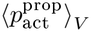 and 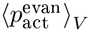, for different activation rates, *k*_act_ and for *δt*_pulse_ = 0.1 s, with *R* = 10 *µ*m, *H* = 4 *µ*m and *R* = 5 *µ*m, *H* = 3 *µ*m.

|  | *R* = 10 μm, *H* = 4 μm |  | *R* = 5 μm, *H* = 3 μm |  |
|---|---|---|---|---|
| *k*<sub>act</sub> (/s) | ⟨*p*<sub>act</sub><sup>prop</sup>⟩<sub>*V*</sub> | ⟨*p*<sub>act</sub><sup>evan</sup>⟩<sub>*V*</sub> | ⟨*p*<sub>act</sub><sup>prop</sup>⟩<sub>*V*</sub> | ⟨*p*<sub>act</sub><sup>evan</sup>⟩<sub>*V*</sub> |
| 10<sup>6</sup> | 0.024 | 0.0021 | 0.083 | 0.0083 |
| 10<sup>7</sup> | 0.14 | 0.015 | 0.4 | 0.057 |
| 10<sup>8</sup> | 0.38 | 0.09 | 0.78 | 0.21 |
| 3×10<sup>8</sup> | 0.47 | 0.18 | 0.9 | 0.46 |

#### Constraints imposed by the width of the spatial distribution of recruited molecules on *k*_act_ and *K*_*b*_

Considering now the width of the spatial distribution of recruited molecules, the value of *k*_act_ should be compatible with the widths measured in different cells (since we assume that *k*_act_ remains unchanged across cells). As references, we chose the narrowest distributions found for batch-1 (freshly cultured cells), anticipating that smaller *K*_*b*_ values would make it possible to fit the broader distributions of batch-2.

In general, the larger *k*_act_ the broader the distribution of recruited molecules gets, due to saturation of photoactivation. Within a certain range, it can be compensated by a larger *K*_*b*_ which leads to more efficient adsorption on the membrane. As a consequence, the width of recruited molecules is relatively stable when *k*_act_ and *K*_*b*_ simultaneously change over two orders of magnitude, as shown in Fig. 8 where the computed recruitment profiles are compared to the narrowest experimental HWHM. However, this compensation effect is no more effective if *K*_*b*_ *>* 10 *µ*m/s. Indeed, using *D* = 1 *µ*m^2^/s, we checked that the shape of the distribution of recruited molecules no longer depends on *K*_*b*_ if it increases beyond 10 *µ*m/s (data not shown). This is because the diffusion length, *λ*_*DK*_ = *D/K*_*b*_, becomes much smaller than the cell dimensions, i.e. the molecules are adsorbed the first time they hit the surface. Consequently, a value *k*_act_ *>* 10^7^ /s inevitably results in theoretical distributions too broad compared to what has been observed for batch-1. Hence the maximum possible value of *k*_act_ is 10^7^ /s, in apparent contradiction with the previous paragraph.

**Figure 8.**
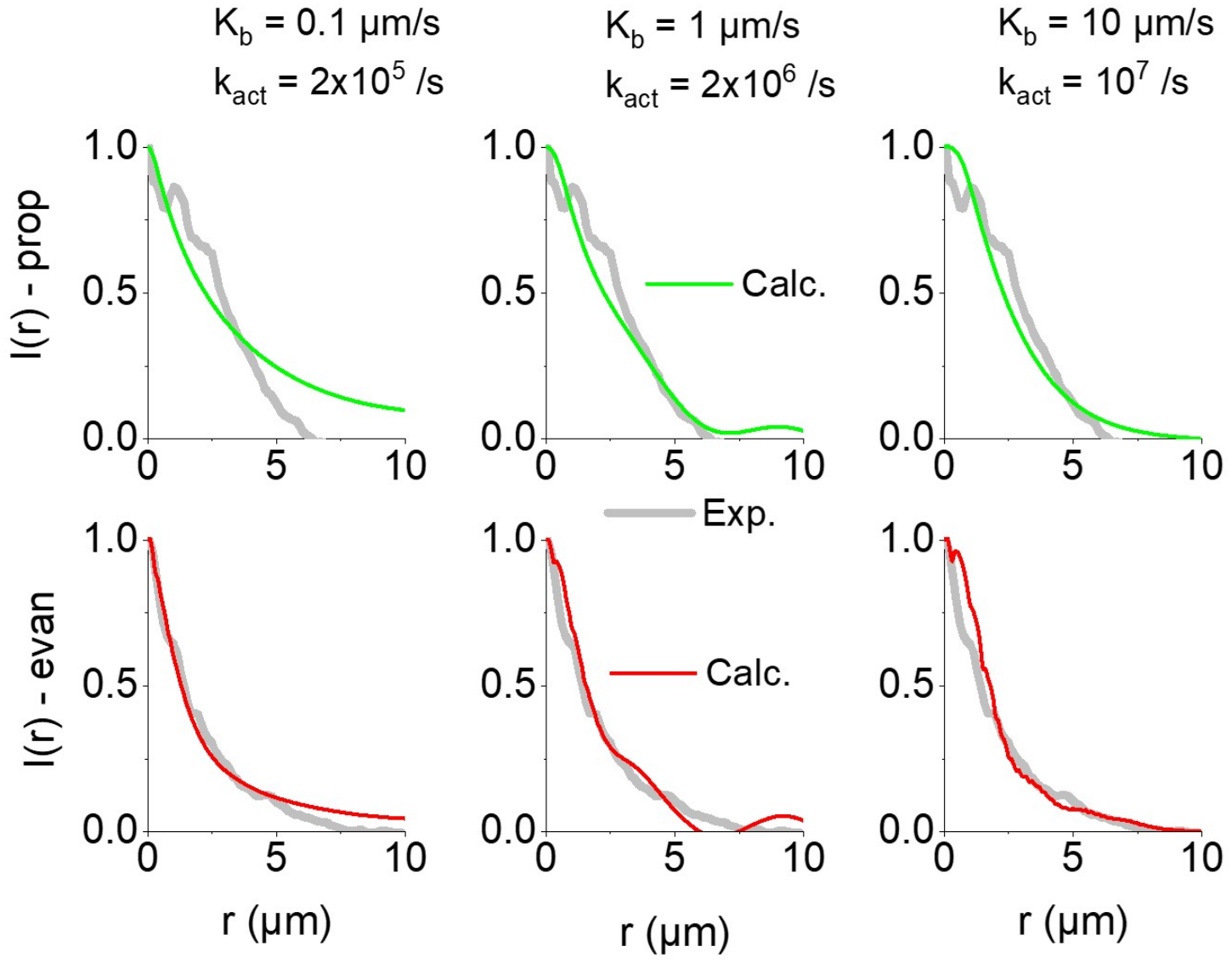
Counterbalancing roles of the photoactivation rate, *k*_act_ and of the surface binding constant, *K*_*b*_, for the calculated recruitment profiles. From left to right *k*_act_ is increased by a factor 50 and *K*_*b*_ by 2 orders of magnitude, while the width of the calculated distribution of recruited molecules stays more or less the same. The thick gray curves correspond to the thinnest distributions of batch-1, observed for the propagative (top) and evanescent (bottom) modes. The pulse duration, *δt*_pulse_, is fixed to 0.1 s while the radius and thickness of the close volume are *R* = 10 *µ*m and *H* = 4 *µ*m.

Going back to Table 2, one can see that the theoretical value of the recruitment slowdown parameter becomes compatible with experiments at *k*_act_ ≈ 10^7^ /s if the dimensions of the cell are reduced to *R* = 5 µm and *H* = 3 µm. Although these values seem much smaller than the physical size of the cells we studied, they should be understood as effective dimensions and include a variety of cellular properties related to compartmentalization (e.g. excluded-volume effects due to organelles and cytoskeletal structures), local heterogeneities in cytoplasmic viscosity or molecular accessibility, molecular crowding effects or hindered diffusion, etc.

#### Final bracketing of *k*_act_ and *K*_*b*_ values

To refine the parameters, the laser pulse duration should also be taken into account. As shown in Table 3, a longer pulse duration increases the theoretical value of the recruitment slowdown parameter, in agreement with the tendency observed in experiments (Table 1). Note that the experiment corresponding to the narrowest distribution has been performed using *δt*_pulse_ = 0.5 s. In Fig. 9, comparisons between the experimental and theoretical distributions are shown. We can conclude that, using *K*_*b*_ = 10 *µ*m s or larger and *k*_act_ = 2 × 10^6^ s or 10^7^ s, the theoretical model is uantitatively compatible with our experimental observations.

**Table 3:**
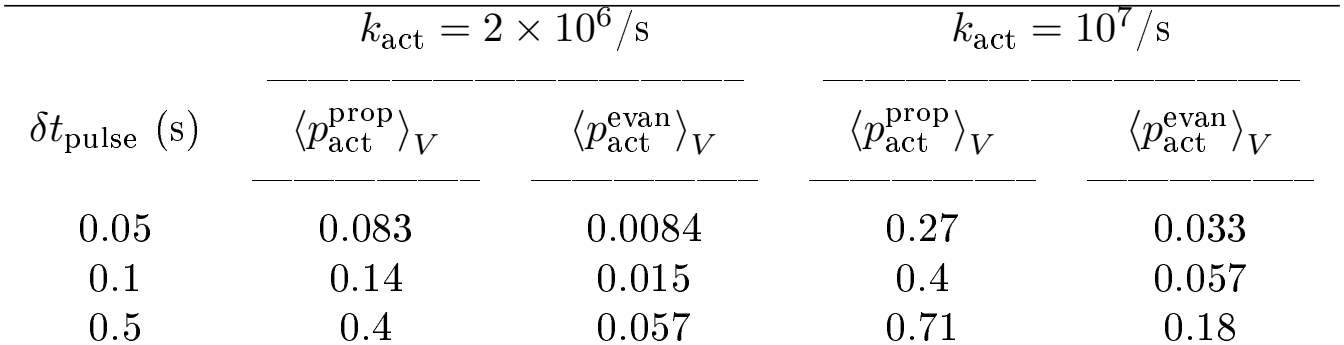
Theoretical values of the slowdown parameters, 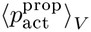 and 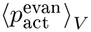, for two activation rates, *k*_act_ = 2 × 10^6^ s and 10^7^ s and for *δt*_pulse_ = 0.05, 0.1 s and 0.5 s. The dimensions are set to *R* = 5 *µ*m and *H* = 3 *µ*m.

|  | k<sub>act</sub> = 2 × 10<sup>6</sup>/s |  | k<sub>act</sub> = 10<sup>7</sup>/s |  |
|---|---|---|---|---|
| δt<sub>pulse</sub> (s) | ⟨p<sub>act</sub><sup>prop</sup>⟩<sub>V</sub> | ⟨p<sub>act</sub><sup>evan</sup>⟩<sub>V</sub> | ⟨p<sub>act</sub><sup>prop</sup>⟩<sub>V</sub> | ⟨p<sub>act</sub><sup>evan</sup>⟩<sub>V</sub> |
| 0.05 | 0.083 | 0.0084 | 0.27 | 0.033 |
| 0.1 | 0.14 | 0.015 | 0.4 | 0.057 |
| 0.5 | 0.4 | 0.057 | 0.71 | 0.18 |

**Figure 9.**
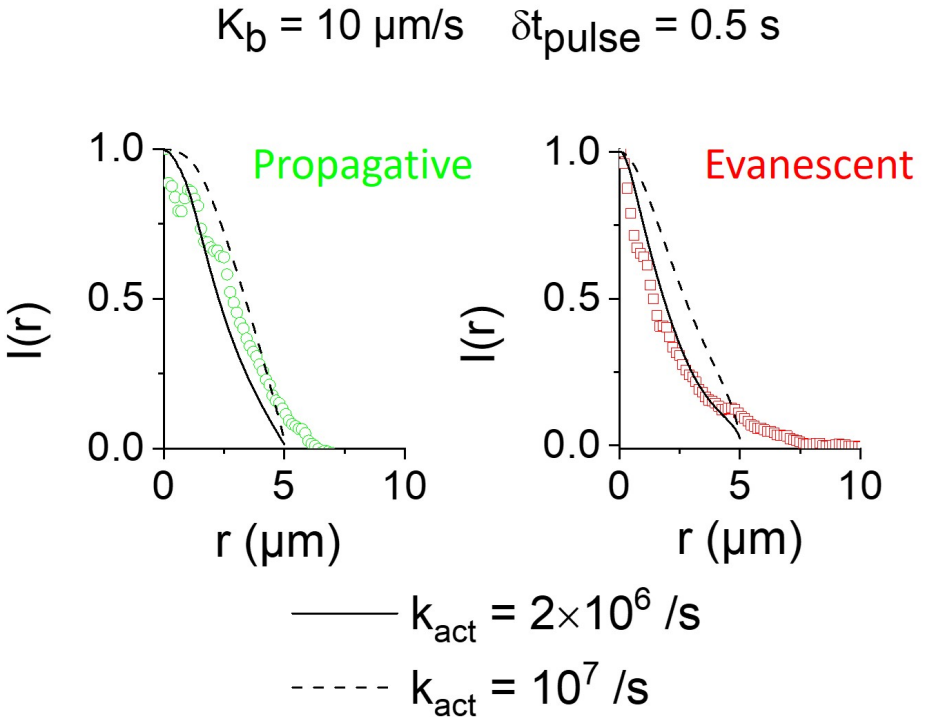
Comparison between the thinnest experimental distributions of batch-1 cells in propagative (left) and evanescent (right) modes and theoretical calculations performed with *K*_*b*_ = 10 *µ*m s, *δt*_p_ = 0.5 s and *k*_act_ = 2 × 10^6^ s or 10^7^ s. The dimensions are set to *R* = 5 *µ*m and *H* = 3 *µ*m.

## Discussion

We have shown above that it is possible to quantitatively describe our optogenetic experiments by using a simple model, where the CRY2 molecules are confined within a closed volume and, once photoactivated by a focused laser spot, either propagative or evanescent, bind forever to the surfaces representing the cellular plasmic membrane. The repetitive laser pulses progressively empty the reservoir of available proteins, inducing a decrease of the absolute number of molecules recruited after each pulse. Both the slowdown of the recruitment rate and the width of the distribution of molecules recruited at the basal surface can be reproduced by the theoretical calculations that depend on a limited number of parameters: those corresponding to the photophysics of the activation process (*σ*_abs_ , *k*_rel_ and *k*_act_) and one quantifying the binding kinetics at the membrane, *K*_*b*_. Using prior knowledge, we fix the value of the diffusion constant, *D* and decided to let vary only the photophysical parameter controlling the transfer from the excited molecular level to the activated state, *k*_act_. Taking into account the whole set of experiments, we can estimate that the photophysical activation rate of CRY2, *k*_act_, falls in the range 2 × 10^6^ s ∼ 10^7^ s. The recruitment width for cells in batch-1 (freshly cultured) indicates that their *K*_*b*_ value ≥ 10 µm.s^*−*1^. Conversely, since the recruitment width of the cells of batch-2 (freshly thawed) is typically twice as large, the corresponding *K*_*b*_ value, and therefore the CIBN membrane concentration, is at least 2 orders of magnitude lower than for freshly cultured cells.

Our experiments were initially planned to study the narrowing of the distribution of molecules recruited after photoactivation by an evanescent laser spot, compared to a propagative one. Contrary to what one might hope, the observed distribution of recruited molecules are much broader than the laser spots themselves and the membrane distribution of molecules recruited with an evanescent spot is only moderately narrower than with a propagative one. Our theoretical model revealed that these relatively large observed distributions are mostly due to the saturation of the photoactivation process, which is inherent to the high irradiance of the laser spot (≈ 1 kW/cm^2^). In that respect, we calculated that a reduction in photoactivation laser power by a factor of 10 and 100 would reduce the distribution widths by a factor of 1.5 and 2.7, respectively (data not shown). However, this would also imply a much smaller number of photoactivated and bound molecules at each pulse, i.e. a much weaker fluorescence signal, making the experiments very difficult to conduct (especially in the presence of cellular autofluorescence). Correspondingly, the photoactivation of the same final number of recruited proteins would require much more time (several tens of min instead of around 1 min), a period of time during which the recruited proteins would start to unbind (the dissociation time is approximately 3 min [13]). Furthermore, the minimum number of bound CRY2 proteins necessary to induce a biological response (e.g. for signalling, etc.) is another key factor that would practically prevent a drastic reduction of the photoactivation intensity. Interestingly, previous works on spatiotemporal control of optogenetics never reported this photoactivation saturation issue [13, 16, 17]. In fact, ignoring saturation is justified when the incoming energy, necessary to activate enough molecules to induce a signal, is diluted over ROIs larger than the laser spot (using either a raster scan or a DMD device).

It is therefore interesting to consider an alternative experimental protocol, compatible with a highly focused laser beam and, as far as we know, never used, where the cell is exposed to the same total amount of light (i.e. same photon budget), but in a continuous way, so that it is illuminated at an irradiance several tens of times weaker than with laser pulses. Doing so, the photoactivation process would be much less saturated, while the time each molecule stay diffusing in the laser beam (0.1 ∼ 1 s) would be comparable to the pulse duration used in the present work. One may thus expect a significant decrease in the distribution width of recruited molecules.

Generally speaking, it is also clear that the development of microscopy platforms and optogenetics to spatiotemporally control cell biological processes and reaction-diffusion systems would benefit from a detailed understanding of the underlying phenomena [26, 27]. This would be the case, for example, of the photosensitive pair, Phy-PIF, which binds in response to red light and dissociates in response to infrared one [18], promising micrometer spatial resolution through a donut-like optogenetic activation.

Further experimental tests of our model should include independent measurements of the membranous concentration of CIBN, as well as recruitment measurements at the apical membrane. Such experiments would help one to assess the contributions of the different membranes to the CRY2-CIBN binding and the impact of the cellular geometry and intracellular compartmentalization on the recruitment rate and the spatial distribution of recruited molecules.

## Conclusion

In this article, we developed a simple, yet non empirical, biophysical model that provides a quantitative description of an optogenetic experiment. It successfully reproduces the observed distribution width of molecules recruited at the basal membrane, as well as their recruitment rate as photoactivation pulses occur over time. This model takes into account the photophysics of the CRY2 molecule, i.e. its excitation and activation by a focused laser beam, providing for the first time an estimation of the intramolecular activation rate of CRY2. The diffusion and interactions of CRY2 molecules at the membrane are described in a non empirical manner, yielding a range of values for the CRY2 binding constant at the membrane, which depends upon the surface density of available CIBN partners. We believe that our theoretical model is solid and general enough to help designing future optogenetic experiments. We put into evidence that, when binding at the membrane is fast enough, the width of the distribution of recruited CRY2 molecules cannot be smaller than a limit depending on diffusion and photoactivation. In this situation, the photophysical saturation is a critical bottleneck that has to be overcome to achieve sub-micrometric resolution.

## Methods

### Microscopy set-up and acquisition protocol

Our setup is based on an inverted microscope stand equipped with a sCMOS camera and has already been described in detail previously [14]. The unique feature of our set-up is the addition of a custom light path including a Digital Micro-mirror Device (DMD) that makes it possible to control which region of the entrance pupil of a high numerical aperture oil objective is illuminated by the incoming laser beam. To generate a laterally confined evanescent field in the sample, only the micro-mirrors of the DMD conjugated with the annular region of the pupil (*n*_oil_ sin(*θ*) *>* 1.33) are in the ON state (i.e. reflects the light towards the objective). Conversely, one obtains a standard propagative laser spot by illuminating the inner region (*n*_oil_ sin(*θ*) *<* 1.33) of the entrance pupil. CRY2 proteins are henceforth activated with a laser beam at 488 nm, focused to be either propagative or evanescent. The laser power at 488 nm was adjusted so that the optical power incident on the back focal plane (i.e. entrance pupil) was the same in both evanescent and propagative modes (P = 4 *µ*W). The fluorescence of mCherry attached to CRY2 proteins is excited at 561 nm and continuously imaged, at the membrane, in Total Internal Reflection Fluorescence (TIRF) configuration.

The acquisition protocol consisted of illuminating each targeted cell by a series of laser pulses at 488 nm, in a propagative or evanescent mode, as shown in Fig. 10A. Then, a 10 min-period in darkness allowed the CRY2-CIBN hetero-dimers to dissociate (unbinding time is ≈ 3 min [13]). Finally, the same cell was submitted to another series of pulses, with similar time parameters, in the other illumination mode. To avoid systematic bias, we varied the order of illumination modes (propagative then evanescent or the inverse order) and also sometimes performed twice the same illumination mode (at the beginning and/or at the end of one cell series of measurements) to compare and verify recruitment images. Experiments have been performed with different numbers of pulses, time intervals between pulses and pulse durations, (*n*_pulse_ , Δ*t*_inter_, *δt*_pulse_): (5, 30 s, 0.5 s), (5, 10 s, 0.5 s), (15, 3 s, 0.1 s), (15, 3 s, 0.05 s), (45, 3 s, 0.1 s). Altogether we collected 30 acquisitions on two batches of cells: one freshly cultured and the other freshly thawed, which are named batch-1 and batch-2 in the following.

**Figure 10.**
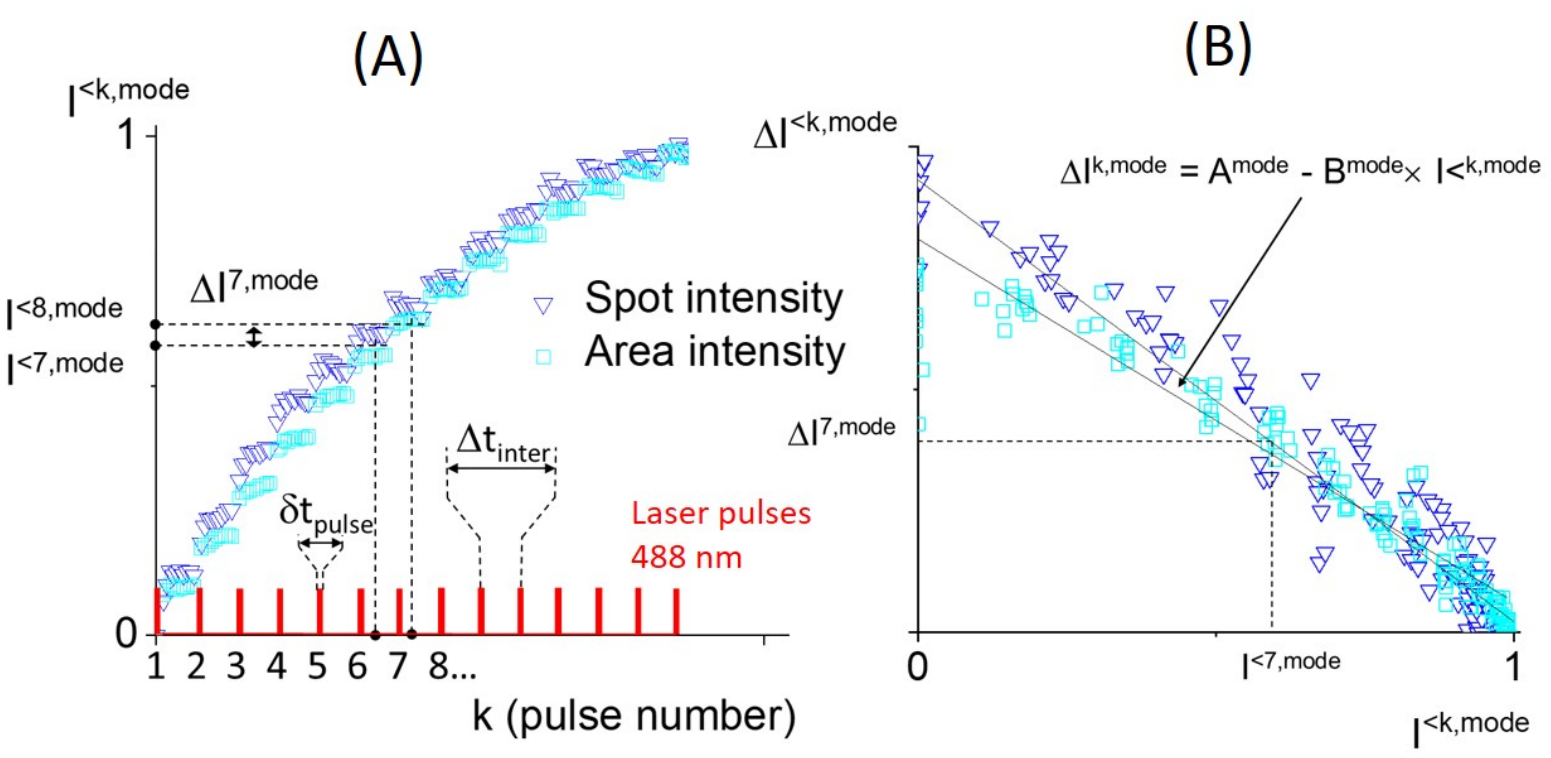
Acquisition protocol and analysis. (A) Each cell is illuminated by a number, *n*_pulse_ , of laser pulses at 488 nm, of duration *δt*_pulse_ and inter-pulse time Δ*t*_inter_. The mCherry fluorescence intensity of the molecules recruited at the membrane, *I*^*<k*, mode^ , is continuously measured simultaneously to the series of pulses, either at the maximum of the TIRF image (triangles for Spot intensity) or by integration over the cell area (squares for Area intensity). (B) The difference between the intensities measured at times separated by Δ*t*_inter_, noted Δ*I*^*k,mode*^, is plotted *versus* the intensity *I*^*k,mode*^, showing a linear decrease.

### Living cell preparation

The experiments have been performed on epithelial Madin-Darby canine kidney (MDCK) cells stably expressing CRY2-mCherry and CIBN-CAAX using a lentiviral transfection [12]. Two batches of cells were considered: freshly cultured cells (batch-1) and freshly thawed ones (batch-2). Further details were given in our previous publication [14].

### Image and data processing

The cells were continuously imaged under 561 nm TIRF excitation, starting approximately 10 s before the series of photoactivation pulses. The images recorded before the very first pulse are averaged, to provide a background that is subtracted to each subsequent image. Then, the mean image intensity within a 1 *µ*m radius disk around the laser spot (Spot intensity) is used to represent the number of molecules bound at the basal membrane just before the *k*^*th*^ pulse, *I*^*<k*,mode^ , as shown in Fig. 10A. Here, the superscript “mode” denotes both the illumination geometry (propagative or evanescent) and the pulse duration, *δt*_pulse_ An alternative estimation of the number of bound molecules is also calculated by integrating the intensity over the cell area (Area intensity). Finally, by angularly averaging the image around the laser spot we obtained a radial profile of the intensity at each time-point, 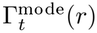.

### Analyses of the molecule recruitment rate and distribution width

#### Analysis of the molecule recruitment rate

We show in the section S1 of Supporting Information, Eq. 41, that the increase induced by the *k*^*th*^ laser pulse, or step in the number of recruited molecules, can be written as a linear function of the number of molecules recruited before the *k*^*th*^ pulse. In terms of intensities, the intensity step, Δ*I*^*k*,mode^ is a linear function of the intensity, *I*^*k*, mode^, before the *k*^*th*^ pulse:

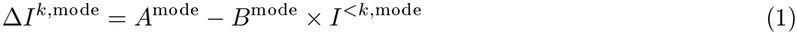

where the step due to the *k*^*th*^ pulse is the difference between the number of molecules before and after the *k*^*th*^ pulse (i.e. before the (*k* + 1)^*th*^ pulse):

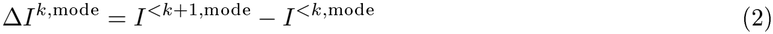

In practice, rather than assigning steps in the recruitment data, we simply paired points separated by the inter-pulse time Δ*t*_inter_ to obtain the values of Δ*I*^*k*,mode^ Indeed, due to continuous time-lapse imaging, a few tens of images are typically recorded per sliding window of duration Δ*t*_inter_. Fig. 10B shows the graph of Δ*I*^*k*, mode^ as a function of *I*^*<k*, mode^ that corresponds to the graph of Fig. 10A.

The negative sign of the second term of the right-hand side of Eq. 1 means that the recruitment rate slows down, which is due to the emptying of the cytosolic reservoir of CRY2 molecules that can be activated by successive activation pulses. Note that the parameter *A*^mode^, which is the recruitment step induced by the very first pulse, depends on the initial concentration of activable CRY2 molecules and cannot be easily controlled (e.g. *A*^mode^ differs for successive acquisitions with the same illumination mode on the same cell). Contrarily, the parameter *B*^mode^ that quantifies the *slowdown of the recruitment rate* is an intensive quantity and can be compared between different acquisitions. In this article, it is also be named *recruitment slowdown parameter*.

#### Analysis of the distribution width

To study the distribution width of the molecules recruited at the basal membrane, we discarded angular information as our theoretical developments are performed with a rotationally symmetric model (see section S1 in Supporting Information). To characterize the width, we used the half width at half maximum, HWHM^mode^ of the radial profile, 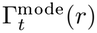, since it should not be much impacted by the wings of the distribution, if at all. As reported in our previous work [14], we have never observed important variation of the distribution shape as the pulses accumulated over time. Consequently, we estimated HWHM^mode^ at the time of maximum recruitment where the signal-to-noise ratio is optimal.

### Statistics

The statistical analyses of experimental outputs have been performed using JASP software [28].

## Associated content

### Supporting information

The Supporting Information is available free of charge at https://pubs.acs.org/doi/…….

Principles of the biophysical model; Photoactivation of CRY2 molecules; Diffusion and surface binding; Surface recruitment of CRY2 molecule (S1 - Theoretical derivation). Sampling methods and software used (S2 - Numerical details).

## Author information

### Corresponding Author

**Irene Wang** - *University Grenoble Alpes, CNRS, LIPhy, 38000 Grenoble, France*

### Authors

**Antoine Delon -** *University Grenoble Alpes, CNRS, LIPhy, 38000 Grenoble, France Marc Grosjean* - *University Grenoble Alpes, CNRS, LIPhy, 38000 Grenoble, France*

**Olivier Destaing** - *Institute for Advanced Biosciences, Research Center INSERM 1209, CNRS UMR 5309, University Grenoble Alpes, France*

**Alexei Grichine** - *Institute for Advanced Biosciences, Research Center INSERM 1209, CNRS UMR 5309, University Grenoble Alpes, France*

## Author Contributions

M.G., I.W. and A.G. conceived and designed the experiments, with inputs from A.D. M.G. run the experiments and acquired the data. O.D. generated all constructs and cell lines, participated to biological validation and data interpretation and provided funding with A.D. A.D. developed the biophysical model, performed the corresponding calculations and analysed the data. A.D. wrote the manuscript with inputs from I.W. and all other authors.

## Acknowledgements

The authors thank M. Balland, T. Boudou, G. Cappello, B. Fourcade and M. Pezet, for stimulating discussions. The authors also thank Philippe Moreau for technical support on the setup, Jean Revilloud for advice on cell culture, Christiane Oddou for molecular biology, Cristina Torres and Lucia Campos-Perello for preparing the cells used in this work.

Agence Nationale de la Recherche (ANR-20-CE42-0018); Fondation ARC pour la Recherche sur le Cancer (ARC 2021 PJA3). The LIPhy and MicroCell core facilities are members of the Imagerie-Sciences du Vivant - In vitro (ISDV) federation, which is labeled by the GIS-IBISA and the France Bioimaging supported by the French National Research Agency (ANR-24-INBS-0005 FBI BIOGEN).

## Notes

### Competing Interest Statement

The authors have declared no competing interest.

