## Supporting Information for "Predictive modeling of photoactivation, diffusion and adsorption in cell The case of the optogenetic CRY2-CIBN system"

**Summary**

Principles of the biophysical model; Photoactivation of CRY2 molecules; Diffusion and surface binding; Surface recruitment of CRY2 molecule (S1 - Theoretical derivation). Sampling methods and software used (S2 - Numerical details).

#### S1 - Theoretical derivation

##### Principles of the biophysical model

In this work, we drew inspiration from the 1D theoretical model of Shvartsman[1] that we extended by representing a cell as a closed volume in the shape of a cylinder of radius $R$, separated by a vertical wall of height $H$, as shown in the main text. The intracellular medium is supposed homogeneous and initially filled with a uniform concentration of CRY2 molecules having a diffusion constant $D$. Once photoactivated during the incoming laser beam pulse, they bind to the surfaces with a uniform trapping rate, $K_b$. Due to the cylindrical geometry, horizontal and vertical molecular movements are uncoupled and the 3D diffusion equation breaks down into a 2D equation and a 1D one, with their respective concentrations, $C_{2D}(r, \theta, t)$ and $C_{1D}(z, t)$.

##### Photoactivation of CRY2 molecules

Each laser pulse of duration $\delta t_{\text{pulse}}$, focused at the interface between the glass substrate and the cellular medium, generates a 3D concentration of photoactivated CRY2 molecules, proportional to the cytosolic concentration -assumed to be homogeneous- of CRY2 molecules that are available just before the $k^{th}$ pulse, $C^{<k,\text{mode}}$:

$$C_{\text{act}}^{k,\text{mode}}(r_0, z_0) = C^{<k,\text{mode}} \times p_{\text{act}}^{\text{mode}}(r_0, z_0) \tag{1}$$

To simplify mathematical expressions, the superscript "mode" stands for both the spatial mode of excitation (propagative *versus* evanescent) and its duration, $\delta t_{\text{pulse}}$. Note that, since the previous pulses have already depleted the cytosolic concentration of available CRY2 molecules, this concentration depends upon the mode and duration of excitation. $p_{\text{act}}^{\text{mode}}(r_0, z_0)$ is the relative population of molecules, located at point $r_0, z_0$ and photoactivated during the time interval $\delta t_{\text{pulse}}$ by the laser beam of rotation symmetry (hence the absence of the variable $\theta_0$). To calculate this population, we assumed that the molecules are immobile during each pulse. This is justified by the small value of the effective diffusion constant, $D \approx 1$ $\mu$m$^2$/s, such that the molecules explore less than 1 $\mu$m in any direction during the illumination time (maximum pulse duration was 0.5 s). This assumption makes it possible to use a simple three-level model, based on photophysical transition rates. More precisely, we consider an excited level with a constant relaxation rate to the ground level, $k_{\rm rel}$ and a constant activation rate to the photoactivated level, $k_{\rm act}$. The excited level is populated by absorption from the ground level with a rate $k_{\rm abs}(r_0, z_0) = \sigma_{\rm abs} \times I^{\rm mode}_{\rm exc}(r_0, z_0)$, where $\sigma_{\rm abs}$ is the absorption cross-section and $I^{\rm mode}_{\rm exc}(r_0, z_0)$ is the excitation intensity of the incoming, propagative or evanescent, laser beam. We ignore any other processes like relaxation from the photoactivated level during and after illumination. The generic solution of this three-level population model based on rate equations can be found in many textbooks such as [2]. In our case it leads to the following solution, assuming that all the CRY2 molecules are initially in their ground level:

$$p^{\rm mode}_{\rm act}(r_0, z_0) = 1 + A \exp(-\nu_{\rm fast}\delta t_{\rm pulse}) - (1 + A) \exp(-\nu_{\rm slow}\delta t_{\rm pulse}) \tag{2}$$

with:

$$\nu_{\rm fast} = \frac{k_{\rm tot}}{2}\left(1 + \sqrt{1 - 4\frac{k_{\rm act}k_{\rm abs}}{k^2_{\rm tot}}}\right) \tag{3}$$

$$\nu_{\rm slow} = \frac{k_{\rm tot}}{2}\left(1 - \sqrt{1 - 4\frac{k_{\rm act}k_{\rm abs}}{k^2_{\rm tot}}}\right) \tag{4}$$

$$k_{\rm tot} = k_{\rm abs} + k_{\rm act} + k_{\rm rel} \tag{5}$$

$$A = \frac{k_{\rm act}k_{\rm abs}}{\nu_{\rm fast}(\nu_{\rm fast} - \nu_{\rm slow})} \tag{6}$$

where the dependencies over $r_0, z_0$, originating from the $k_{\rm abs}$ term, have been skipped over for sake of simplicity.

The excitation intensity, $I^{\rm mode}_{\rm exc}(r_0, z_0)$, has been calculated by considering an incident planar scalar wave focused, at the glass - cellular medium interface, with an oil objective of numerical aperture NA $= 1.49$. The calculation of $I^{\rm mode}_{\rm exc}$ was performed using the angular spectrum method, similarly to that described in our previous article [3]. The propagative configuration corresponds to a pupil support given by a disk of external radius $R_{disk} = 1.3 f_{obj}$, $f_{obj}$ being the objective image focal distance. Conversely, when the pupil support is a ring of inner and outer radii $R_{ring,in} = 1.36 f_{obj}$ and $R_{ring,out} = 1.49 f_{obj}$ respectively, an evanescent beam is generated in the aqueous medium beyond the cover-slip surface. Using Axelrod's theory [4], we found that the absorption probability of photons just above the interface is twice higher in evanescent mode compared to the propagative one, which is a consequence of the intensity peak when the angle of the incident wave equals the critical angle. Given the laser power falling on the back focal plane (or entrance pupil) of the objective, this leads to an equivalent power at the interface of $10^{13}$ and $2 \times 10^{13}$ photons/s in propagative and evanescent mode, respectively. Compared to the propagative case, the striking feature of a focused evanescent beam is, in addition to the well known axially exponentially decreasing intensity, the presence of lateral side lobes around the center [5]. Finally, to mimic our experimental situation where we observed optical aberrations [3], the incoming wavefront has been imparted with a random superimposition of 25 Zernike modes $Z^m_n$, where the $n$ index ranges from 2 to 6. The amplitudes of the Zernike modes follow a centered gaussian distribution, the rms of which has been chosen to reproduce the observed lateral profiles shown in Fig. 4 of Ref. [3] (rms = 2.3 rad for the propagative mode and 1.74 rad for the evanescent one). Practically, we have generated 20 random superimpositions of Zernike modes, the corresponding excitation intensities of which, $I^{\rm mode}_{\rm exc}$, was eventually angularly averaged to fulfill the radial symmetry of our analytical model.

#### Diffusion and surface binding

##### Horizontal diffusion

The concentration in the horizontal plane obeys to the diffusion equation that is, in polar coordinates:

$$\frac{\partial C_{2D}}{\partial t} = D(\frac{1}{r}\frac{\partial}{\partial r}r\frac{\partial}{\partial r} + \frac{1}{r^2}\frac{\partial^2}{\partial\theta^2})C_{2D} \tag{7}$$

and to the boundary conditions at the lateral wall, including the Fick law and the trapping rate:

$$j_r(R,\theta,t) = -D\frac{\partial C_{2D}}{\partial r}\bigg|_{r=R} = K_b C_{2D}(R,\theta,t) \tag{8}$$

where $j_r$ stands for the radial 2D flow. Eq. 7 being separable *versus* the $r, \theta$ and $t$ variables, the general 2D solution can be written as the following discrete series:

$$C_{2D}(r,\theta,t) = \sum_l \sum_m [w^c_{l,m} A_l \cos(l\theta) + w^s_{l,m} B_l \sin(l\theta)] C_{l,m} J_l(k_{l,m} r) e^{-Dk_{l,m}{}^2 t} \tag{9}$$

where $J_l$ are the $1^{st}$ kind Bessel functions of integer order $l$ ($l \geq 0$). $w^c_{l,m}$ and $w^s_{l,m}$ are constants that depend upon the initial condition, while $A_l, B_l$ and $C_{l,m}$ are normalisation constants. The later are chosen so to normalise the angular and radial basis functions, which leads to:

$$A_0 = \frac{1}{\sqrt{2\pi}} \qquad B_0 = 0 \tag{10}$$

$$l > 0 : A_l = \frac{1}{\sqrt{\pi}} \qquad B_l = \frac{1}{\sqrt{\pi}} \tag{11}$$

$$C_{l,m} = \frac{1}{\sqrt{\int_0^R r J_l^2(k_{l,m} r) dr}} \tag{12}$$

The $k_{l,m}$ values are obtained by inserting the general 2D solution, Eq. 9, in the boundary condition, Eq. 8 and solving it:

$$K_b J_l(k_{l,m} R) = -D\frac{\partial J_l(k_{l,m} R)}{\partial R} \tag{13}$$

Note that $k_{l,m} = 0$ is not a physical solution of this later equation, except when $K_b = 0$, a situation that we do consider in this work (since it would correspond to no adsorption at the surfaces). We discuss, in the numerical detail section, the maximum value of $l$ and the number of $k_{l,m}$ solutions that are necessary for a proper representation of the 2D concentration.

We are now left to determine the $w^c_{l,m}$ and $w^s_{l,m}$ weights in the series of Eq. 9, which depend upon the initial condition. For a molecule initially located at $r_0, \theta_0$, the 2D concentration at time $t = 0$, $C_{2D}(r,\theta,0;r_0,\theta_0)$, is given by $\frac{1}{r}\delta(r-r_0)\delta(\theta-\theta_0)$. Thanks to the Sturm–Liouville theory, the $A_l\cos(l\theta)C_{l,m} \times J_l(k_{l,m}r)$ and $B_l\sin(l\theta)]C_{l,m}J_l(k_{l,m}r)$ functions form an orthogonal basis, according to which:

$$w^c_{l,m} = \int_0^R \int_0^{2\pi} C_{2D}(r,\theta,0;r_0,\theta_0) A_l \cos(l\theta) C_{l,m} J_l(k_{l,m} r) r dr d\theta \tag{14}$$

$$w^s_{l,m} = \int_0^R \int_0^{2\pi} C_{2D}(r,\theta,0;r_0,\theta_0) A_l \sin(l\theta) C_{l,m} J_l(k_{l,m} r) r dr d\theta \tag{15}$$

One thus obtains:

$$w^c_{l,m} = A_l \cos(l\theta_0) C_{l,m} J_l(k_{l,m} r_0) \tag{16}$$

$$w^s_{l,m} = B_l \sin(l\theta_0) C_{l,m} J_l(k_{l,m} r_0) \tag{17}$$

Therefore the general 2D solution for an initial position at $r_0, \theta_0$ is simply written as follows:

$$C_{2D}(r,\theta,t;r_0,\theta_0) = \sum_l \sum_m [A_l^2 \cos(l\theta_0)\cos(l\theta) + B_l^2 \sin(l\theta_0)\sin(l\theta)] \times C^2_{l,m} J_l(k_{l,m} r_0) J_l(k_{l,m} r) e^{-Dk_{l,m}{}^2 t} \tag{18}$$

**Vertical diffusion**

The concentration along the vertical z axis obeys to the diffusion equation:

$$\frac{\partial C_{1D}}{\partial t} = D\frac{\partial^2 C_{1D}}{\partial z^2} \tag{19}$$

and to the boundary conditions at the basal and apical surfaces, including the Fick law and the trapping rate:

$$j_z(0,t) = -D\frac{\partial C_{1D}}{\partial z}\bigg|_{z=0} = -K_b C_{1D}(0,t) \tag{20}$$

$$j_z(H,t) = -D\frac{\partial C_{1D}}{\partial z}\bigg|_{z=H} = K_b C_{1D}(H,t) \tag{21}$$

where $j_z$ stands for the 1D flow. Next, since Eq. 19 is separable *versus* the $z$ and $t$ variables, the general 1D solution can be written as the following discrete series:

$$C_{1D}(z,t) = \sum_n w_n c_n(z) e^{-Dk_n{}^2 t} \tag{22}$$

with $w_n$ the weight of the $n$ component in the series representing the 1D concentration and

$$c_n(z) = A_n \cos(k_n z) + B_n \sin(k_n z) \tag{23}$$

The possible values of $k_n$ appearing in Eq. 22,23 are obtained by combining these equations with the two boundary conditions. The first one at $z = 0$, Eq. 20, implies that $c_n(z)$ must read:

$$c_n(z) = B_n \left[\frac{Dk_n}{K_b}\cos(k_n z) + \sin(k_n z)\right] \tag{24}$$

while the $B_n$ values are determined by normalizing the basis functions, $c_n(z)$:

$$\int_0^H c_n(z)^2 dz = 1 \tag{25}$$

Using the second boundary condition, Eq. 21 , one obtains an equation whose solution gives the discrete and positive values $k_n$:

$$\cot k_n H = \frac{1}{2}\left(\frac{Dk_n}{K_b} - \frac{K_b}{Dk_n}\right) \tag{26}$$

We discuss, in the numerical detail section, the number of $k_n$ solutions that is adapted to the present situation.

We now consider the case of a molecule initially located at $z_0$, i.e. $C_{1D}(z,0;z_0) = \delta(z - z_0)$. The weight, $w_n$, of each component in the series of Eq. 22 is determined by using the fact that, according to the Sturm–Liouville theory, the $c_n(z)$ functions form an orthogonal basis:

$$w_n = \int_0^H C_{1D}(z,0;z_0) c_n(z) dz \tag{27}$$

$$= c_n(z_0) \tag{28}$$

Therefore the general 1D solution for an initial position at $z_0$ is:

$$C_{1D}(z,t;z_0) = \sum_n c_n(z_0) c_n(z) e^{-Dk_n{}^2 t} \tag{29}$$

In the numerical detail section we discuss the maximum value of $n$ required to properly represent the 1D concentration.

##### 3D diffusion and binding to the surfaces

By combining the solutions of the horizontal and vertical diffusion equations one obtains the solution for the 3D diffusion from an initial point:

$$C_{3D}(r,\theta,z,t;r_0,\theta_0,z_0) = C_{2D}(r,\theta,t;r_0,\theta_0)C_{1D}(z,t;z_0) \tag{30}$$

that is:

$$C_{3D}(r,\theta,z,t;r_0,\theta_0,z_0) = \sum_l \sum_m \sum_n e^{-D({k_{l,m}}^2+{k_n}^2)t} c_n(z_0)c_n(z)\times \\ [A_l^2\cos(l\theta_0)\cos(l\theta)+B_l^2\sin(l\theta_0)\sin(l\theta)]C_{l,m}^2 J_l(k_{l,m}r_0)J_l(k_{l,m}r) \tag{31}$$

We are now left to calculate, for an initial position at $r_0,\theta_0,z_0$, the final surface density at any point $r_s,\theta_s,z_s$ of the cylinder surfaces, once the whole pool of proteins has been removed from the volume. This is obtained by integrating the molecular flow at the surfaces, from $t=0$ to $\infty$:

$$\rho_S(r_s,\theta_s,z_s;r_0,\theta_0,z_0) = \int_0^\infty K_b C_{3D}(r_s,\theta_s,z_s,t;r_0,\theta_0,z_0)dt \tag{32}$$

which straightforwardly leads to:

$$\rho_S(r_s,\theta_s,z_s;r_0,\theta_0,z_0) = \sum_l \sum_m \sum_n \frac{K_b}{D({k_{l,m}}^2+{k_n}^2)} c_n(z_0)c_n(z_s)\times \\ [A_l^2\cos(l\theta_0)\cos(l\theta_s)+B_l^2\sin(l\theta_0)\sin(l\theta_s)]C_{l,m}^2 J_l(k_{l,m}r_0)J_l(k_{l,m}r_s) \tag{33}$$

where $z_s=0$ and $H$ for the basal and apical surfaces, respectively. An equivalent solution would be obtained for proteins binding to the lateral wall, in which case $r_s=R$.

A consequence of the present model is that two important characteristic lengths drive the recruitment of molecules at the surfaces: the distance between the surfaces, $H$ and the diffusion length, $\lambda_{DK}=D/K_b$. $\lambda_{DK}$ is the characteristic distance over which the diffusion and reaction flows balance each other. If $\lambda_{DK}\gg H$ the concentration along the $z$ axis tends to uniformity, so that the molecules are equally adsorbed on both surfaces (they diffuse back and forth between the walls before being adsorbed). Conversely, when $\lambda_{DK}\leq H$, molecules are mainly recruited at the surface they are closest to.

#### Surface recruitment of CRY2 molecules

##### Spatial distribution of recruited proteins

Each laser pulse is followed by the recruitment of all the proteins activated in the cylindric volume, that is by an overall increase of the surface density of molecules. This increase, induced by the $k^{th}$ pulse of duration $\delta t_{\text{pulse}}$, is thus calculated by integrating $C_{\text{act}}^{k,\text{mode}}\times\rho_S$ over $r_0,\theta_0,z_0$ which, using Eq. 1, can be written:

$$\Delta\Gamma_S^{k,\text{mode}}(r_s,\theta_s,z_s) = C^{<k}\int_0^H\int_0^R\int_0^{2\pi} p_{\text{act}}^{\text{mode}}(r_0,z_0)\times\rho_S(r_s,\theta_s,z_s;r_0,\theta_0,z_0)dr_0d\theta_0dz_0 \tag{34}$$

Note that the CRY2 cytosolic concentration before the $k^{th}$ pulse, $C^{<k}$, depends upon the excitation mode and its duration but, to simplify the notation, we skip the superscript "mode". The integration over $\theta_0$ disappears due to the rotation symmetry of the relative population of activated molecules, $p_{\text{act}}^{\text{mode}}$. As a matter of fact, the integration of terms $A_l^2\cos(l\theta_0)$ and $B_l^2\sin(l\theta_0)$ appearing in the sum of Eq. 33, equates 0 except for the term $A_0$, leading to:

$$\Delta\Gamma_S^{k,\text{mode}}(r_s,z_s) = C^{<k}\frac{K_b}{D}\sum_m C_{0,m}^2 J_0(k_{0,m}r_s)\sum_n \frac{c_n(z_s)}{k_{0,m}^2+{k_n}^2}\times \int_0^H\int_0^R p_{\text{act}}^{\text{mode}}(r_0,z_0)J_0(k_{0,m}r_0)c_n(z_0)dr_0dz_0 \quad (35)$$

The spatial distribution of proteins recruited as the result of the $k^{th}$ pulse is obtained by setting $z_s$ to 0 or $H$ for the basal or apical surfaces, respectively and $r_s$ to $R$ for the lateral wall.

##### Recruitment rate

We now want to calculate the recruitment rate over the successive photoactivation pulses. As the time interval between two successive laser pulses is at least 3 s, we can assume that the cytosolic concentration of CRY2 molecules capable of being activated homogenises between each pulse, given the micrometric cell dimensions and $D = 1$ $\mu\text{m}^2/\text{s}$. This makes it possible to use Eq. 35 according to which the shape of the surface distribution of recruited molecules does not change over time. To proceed further, we write that the proteins recruited on all the surfaces (i.e. the basal, apical surfaces and the lateral wall) have been formerly photoactivated in the cell volume and never unbind on the time scale of the experiment, that is:

$$\int_S \Delta\Gamma_S^{k,\text{mode}}(\vec{s})d\vec{s} = \int_V C^{<k}p_{\text{act}}^{\text{mode}}(r_0,z_0)2\pi r_0 dr_0 dz_0 \quad (36)$$

$$= C^{<k}V\langle p_{\text{act}}^{\text{mode}}\rangle_V \quad (37)$$

where $\vec{s}$ is the generalised coordinate over all the surfaces $S$ and $V = \pi R^2 H$ is the total volume of the cell. $\langle p_{\text{act}}^{\text{mode}}\rangle_V$ is the mean value, over the volume of the cell, of the relative population of proteins activated under a specific illumination mode (evanescent or propagative). Moreover, the homogeneous concentration of CRY2 molecules that can be activated by the $k^{th}$ pulse, $C^{<k,\text{mode}}$, is the initial concentration at the beginning of the experiment, $C^0$, minus that due to the molecules already bound to the surfaces before that pulse, formally expressed as $V\Delta C_S^{<k,\text{mode}}$, that is:

$$C^{<k,\text{mode}} = C^0 - \Delta C_S^{<k,\text{mode}} \quad (38)$$

with

$$V\Delta C_S^{<k,\text{mode}} = \int_S \Gamma_S^{<k,\text{mode}}(\vec{s})d\vec{s} \quad (39)$$

where $\Gamma_S^{<k,\text{mode}}$ is the spatial distribution of the proteins already bound before the $k^{th}$ pulse. Combining Eq. 36 to 39 one gets:

$$\int_S \Delta\Gamma_S^{k,\text{mode}}(\vec{s})d\vec{s} = \langle p_{\text{act}}^{\text{mode}}\rangle_V\left[VC^0 - \int_S \Gamma_S^{<k,\text{mode}}(\vec{s})d\vec{s}\right] \quad (40)$$

Then, rewriting $\langle p_{\text{act}}^{\text{mode}}\rangle_V VC^0$ as $\int_S \Delta\Gamma_S^{1,\text{mode}}(\vec{s})d\vec{s}$, we are ultimately left with an equation involving the integrands:

$$\Delta\Gamma_S^{k,\text{mode}}(\vec{s}) = \Delta\Gamma_S^{1,\text{mode}}(\vec{s}) - \langle p_{\text{act}}^{\text{mode}}\rangle_V\Gamma_S^{<k,\text{mode}}(\vec{s}) \quad (41)$$

Note that this equation is valid for any of the surfaces (basal, apical or lateral) enclosing the volume of the cell. It means that the recruitment rate following each activation pulse decreases proportionally to the proteins already recruited, with a factor given by the activation efficiency averaged over all the volume. Interestingly, this slowdown parameter, $\langle p_{\text{act}}^{\text{mode}}\rangle_V$, is independent from the diffusion and interaction processes. Note that Eq. 41 is strictly equivalent to Eq. (3) of the L. Valon paper [6] when CRY2 unbinding from the membrane is neglected. The proportion of proteins recruited after a laser pulse, noted $f$ in their paper, corresponds to $\langle p_{\text{act}}^{\text{mode}}\rangle_V$ in the present one.

### S2 - Numerical details

#### Sampling and software used

The spatial distribution of the photoactivation laser beam intensity is calculated using MATLAB [7] and sampled with a radial step of 0.02 $\mu$m and an axial one (along z) of 0.0625 $\mu$m.

Mathematica (Wolfram) [8] has been used, both for cell image analysis and also to calculate the theoretical spatial distributions of recruited molecules. The radial resolution used for the distribution of molecules recruited in the basal plane is 0.1 $\mu$m. The maximum value of the radial wave vector, $k_{l,m}$, is set to $10\pi$ $\mu$m$^{-1}$. The maximum value of the axial wave vector, $k_n$, is $200\pi$ $\mu$m$^{-1}$ to allow calculations with $K_b$ values larger than 10 $\mu$m/s (i.e. suited for a diffusion length, $\lambda_{DK}$, down to 0.01 $\mu$m).

### Author information


#### Corresponding Author

**Irène Wang** - *University Grenoble Alpes, CNRS, LIPhy, 38000 Grenoble, France*

#### Authors

**Antoine Delon** - *University Grenoble Alpes, CNRS, LIPhy, 38000 Grenoble, France*
**Marc Grosjean** - *University Grenoble Alpes, CNRS, LIPhy, 38000 Grenoble, France*
**Olivier Destaing** - *Institute for Advanced Biosciences, Research Center INSERM 1209, CNRS UMR 5309, University Grenoble Alpes, France*
**Alexei Grichine** - *Institute for Advanced Biosciences, Research Center INSERM 1209, CNRS UMR 5309, University Grenoble Alpes, France*


#### Author Contributions

M.G., I.W. and A.G. conceived and designed the experiments, with inputs from A.D. M.G. run the experiments and acquired the data. O.D. generated all constructs and cell lines, participated to biological validation and data interpretation and provided funding with A.D. A.D. developed the biophysical model, performed the corresponding calculations and analysed the data. A.D. wrote the manuscript with inputs from I.W. and all other authors.

### Acknowledgements


The authors thank M. Balland, T. Boudou, G. Cappello, B. Fourcade and M. Pezet, for stimulating discussions.

Agence Nationale de la Recherche (ANR-20-CE42-0018); Fondation ARC pour la Recherche sur le Cancer (ARC 2021 PJA3). The LIPhy and MicroCell core facilities are members of the Imagerie-Sciences du Vivant – In vitro (ISDV) federation, which is labeled by the GIS-IBISA and the France Bioimaging supported by the French National Research Agency (ANR-24-INBS-0005 FBI BIOGEN).

(7) Inc., T. M. MATLAB version: 9.10.0 (R2021a), Natick, Massachusetts, United States, 2021.

(8) Inc., W. R. Mathematica, Version 12.3, Champaign, IL, 2021.
